# Novel eDNA approaches to monitor Western honey bee (*Apis mellifera*) microbial and arthropod communities

**DOI:** 10.1101/2022.08.31.506105

**Authors:** Leigh Boardman, Jose A.P. Marcelino, Rafael E. Valentin, Humberto Boncristiani, Jennifer Standley, James D. Ellis

## Abstract

Pests and pathogens are a continuous threat to the health of Western honey bees *Apis mellifera* L. Monitoring honey bee colonies for arthropod pests, disease-causing bacteria and fungi, and early detection of new invasions is essential to maintain the pollination services provided by honey bees. Here, we investigated the feasibility of using eDNA metabarcoding to detect honey bee pests and pathogens in their hives and across their foraging environment. We sampled 13 different surfaces within and outside hives from our test apiary to determine where the most informative eDNA could be obtained. Furthermore, we compared two main eDNA collection techniques – wiping surfaces with moistened forensic swabs and using a spray/wash technique that washed surface DNA into a bucket, before collecting the DNA on a filter. We used DNA metabarcoding with universal primer sets to target arthropod, bacterial and fungal communities. Our results showed that most surfaces yielded usable DNA, and that results of the swabs and spray/wash methods were similar when they could be applied to the same surface. We were able to detect DNA from the small hive beetle (*Aethina tumida), Varroa destructor* mites and European foulbrood (*Melissococcus plutonius*), all of which matched our visual observations of clinical signs of these pests and pathogens in the hives we tested. DNA from some species was location specific, which has implications for using eDNA as a monitoring tool. Collectively, our data show that eDNA metabarcoding can accurately detect DNA from arthropods and microbes honey bees contact and has the potential to provide information on disease and pest incidence, *Apis* species identity, and *A. mellifera* subspecies identity of the colony and/or apiary. In sum, eDNA metabarcoding can be used as a comprehensive molecular predictor tool for colony health surveys.

## 1. Introduction

Healthy Western honey bee (*Apis mellifera* Linnaeus, 1758) colonies are essential for food security, contributing an estimated $12-50 billion USD/year in pollination services to the U.S. economy (Bauer & Wing, 2010; Calderone, 2012), and $182 - 577 billion USD/year globally (Gallai et al., 2009; Lautenbach et al., 2012). Unfortunately, these services are at risk from many stressors that have led to high gross loss rates of managed honey bee colonies in some areas of the world (Lee et al., 2015). Among these stressors, pests, diseases, and the emergence of novel issues remain a continuous threat to honey bee colony health and productivity (Goulson et al., 2015; Ray et al., 2020). Establishing effective monitoring protocols for these stressors would improve colony health and help ensure the pollination services provided by honey bees. Reliable, easy to administer detection tools would allow apiary inspectors to monitor for potential cryptic threats in a timely manner, thereby safeguarding the health and productivity of managed honey bee colonies before sustaining extensive economic and/or colony losses.

Current methods for monitoring honey bee pests and pathogens are expensive and time-consuming to use. Furthermore, they focus on examining targeted species, thus potentially missing new or emerging pests/pathogens. In the U.S., scientists administering the APHIS National Honey Bee Survey currently monitor for nine honey bee viruses, *Nosema* spp. spores, and *Varroa destructor* (Anderson & Trueman, 2000) loads. Additionally, they confirm the absence of *Tropilaelaps* spp., *Apis cerana* (Fabricius, 1793), and Slow Bee Paralysis Virus (Fahey et al., 2018, 2019; LeBrun, n.d.; Ray et al., 2020; Traynor et al., 2016). Beekeepers also screen their bees visually for American foulbrood (*Paenibacillus larvae*), European foulbrood (*Melissococcus plutonius*), Sacbrood virus, small hive beetles (*Aethina tumida*), and wax moths (*Galleria mellonella* and *Achroia grisella*), among other pests/pathogens.

Current methodologies for monitoring biotic colony threats involve the collection of a significant numbers of individual honey bees and/or their brood. Moreover, the different protocols used require different skillsets, making the process time consuming to execute, costly by necessitating trained personal, and inefficient as it includes only known pathogens and pests, thus restricting the ability of surveys to detect new or emergent threats. There is a critical need to design improved, non-targeted methods to detect honey bee pests and pathogens. The development of such methods could allow one to detect new invasive arthropods and pathogens early, ultimately providing a strong advantage to beekeepers in the constant, ever-evolving battle to keep honey bee colonies healthy.

Environmental DNA (eDNA) is DNA collected in intracellular or extracellular form from environmental samples rather than from biological organisms themselves. When paired with a high-throughput sequencing method like metabarcoding, eDNA offers an indirect and non-targeted identification method that can be used for several applications (Beng & Corlett, 2020; Garlapati et al., 2019; Ruppert et al., 2019). Theoretically, eDNA metabarcoding could be used to detect biotic stressors of honey bees and may serve as a viable detection method to improve current honey bee colony monitoring and diagnostic efforts. High-throughput molecular methods have been used in honey bee research to detect microbes (Cox-Foster et al., 2007) and viruses (Beaurepaire et al., 2020; Galbraith et al., 2018; Granberg et al., 2013; Runckel et al., 2011) in honey bees. eDNA has been used successfully to detect brown marmorated stink bugs in agricultural fields (Valentin et al., 2018), spotted lanternflies in forests (Valentin et al., 2020), and arthropod diversity from house dust (Madden et al., 2016), wild flowers (Thomsen & Sigsgaard, 2019), honeydew honey (Utzeri et al., 2018), and pollinator communities (Harper et al., 2021). Emerging eDNA methods have been used to sample airborne DNA to detect terrestrial insects (Roger et al., 2022), demonstrating the sensitivity of this technique. In several cases, eDNA metabarcoding can match, or exceed, traditional identification and detection methods in a variety of environments (Allen et al., 2021; Bush et al., 2020).

When paired, eDNA and high-throughput molecular methods offer an opportunity for exciting honey bee research. To date, eDNA studies on honey bees have focused on honey. Bovo et al. (2018) used a shotgun metagenomics approach to study two sources of European honey. Together with DNA from host *Apis* sp., they detected *G. mellonela* (Linnaeus, 1758, the greater wax moth), *V. destructor, N. ceranae* (Fries et. al., 1996), *Bettsia alvei* (Betts, Skou, 1972, mold on pollen stored in a hive), *Aspergillus* spp. (fungi that cause stonebrood in honey bee colonies), *M. plutonius* (Truper & Clari, 1998, bacterial agent of European foulbrood), and *Apis mellifera* filamentous virus (AmFV), along with other arthropods, plants, fungi and bacteria. Subsequent studies have used eDNA from honey for *V. destructor* monitoring and biogeography (Utzeri et al., 2019), the entomological authentication of European honey from different *A. mellifera* lineages (Soares et al., 2019), to detect pathogens, parasites, and pests in honey (Ribani et al., 2020, 2021, 2022), and identify organisms in contact with bees while foraging (Bovo et al., 2020).

Here, we aimed to investigate the feasibility of using eDNA metabarcoding to detect honey bee pests and pathogens in their hives and across their foraging environment by determining 1) what substrates or mediums within hives from our testing apiary and immediate surroundings contain the most informative eDNA, 2) what eDNA sampling methods are most effective at collecting informative eDNA, and 3) what arthropod pests, bacteria and fungi are detectable using these methods. Our preference is for finding sampling methods that can be adopted easily by beekeepers themselves or apiary inspectors with minimal training.

## 2. Materials and Methods

All eDNA samples were collected at the University of Florida Honey Bee Research and Extension Laboratory (HBREL), Entomology and Nematology Department, in Gainesville, Florida, USA between 13 and 28 September 2021. This laboratory houses approximately fifty full size, Langstroth-style hives occupied by mixed stock honey bees of European descent. The hives were managed using standard best management practices (feeding when necessary, disease/pest control, swarm control, etc.) for the region. Furthermore, they were inspected for the presence of European foulbrood, small hive beetles, *V. destructor* and greater wax moths by a trained beekeeper to ground truth the eDNA data.

### 2.1 Sampling

Thirteen different samples were collected (Table 1, Table S1, Figure 1). eDNA collection methods consisted of sampling surfaces in the apiary, hives, and foraging paths of honey bees with swabs and spray/wash surface aggregation methods (Valentin et al., 2020). Liquid and solid samples were also collected. Details of each of these methods follow. Three independent replicates were obtained for each sample and method, except for the hive bottom board. Sampling hive bottom boards requires lifting the entire hive, which is disruptive to the colony. Three hive bottom boards were sampled by swabbing, then by the spray/wash surface aggregation method on different parts of each board (details follow). See Table S1 for more information about effort per sample type.

**Table 1.** List of surfaces, liquids, and solids sampled from apiary environments. Numbering refers to Figure 1.

| # | Abbreviation | Sample | Sample Collection Method |  |  |  |
| --- | --- | --- | --- | --- | --- | --- |
|  |  |  | Swab | Roller | Spray / Wash | Direct collection |
| 1 | HBB | Hive bottom board | Yes<sup>1</sup> |  | Yes<sup>1</sup> |  |
| 2 | HER | Hive entrance reducer | Yes |  | Yes |  |
| 3 | HDT | Hive detritus from tray | Yes |  | Yes |  |
| 4 | HT | Hive tools | Yes |  | Yes |  |
| 5 | FF | Frame of foundation (blanked) | Yes | Yes | Yes |  |
| 6 | FDC | Frame of drawn comb (not blanked) | Yes |  | Yes |  |
| 7 | HF | Hive feeder (1:1, water:fructose) |  |  |  | Yes |
| 8 | PT | Plant transect in bee foraging path | Yes |  | Yes |  |
| 9 | PL | Plant litter at base of hive |  |  | Yes |  |
| 10 | SUH | Soil underneath hive |  |  |  | Yes |
| 11 | PW | Pond water |  |  |  | Yes |
| 12 | SP | Soil next to pond<sup>2</sup> |  |  |  | Yes |
| 13 | RJ | Royal jelly |  |  |  | Yes |

<sup>1</sup> These samples are not independent. Data were combined as swabs and spray were performed on the same hive bottom board.
<sup>2</sup> Soil next to pond failed to pass bioinformatic filtering across all four datasets.

**Figure 1.**
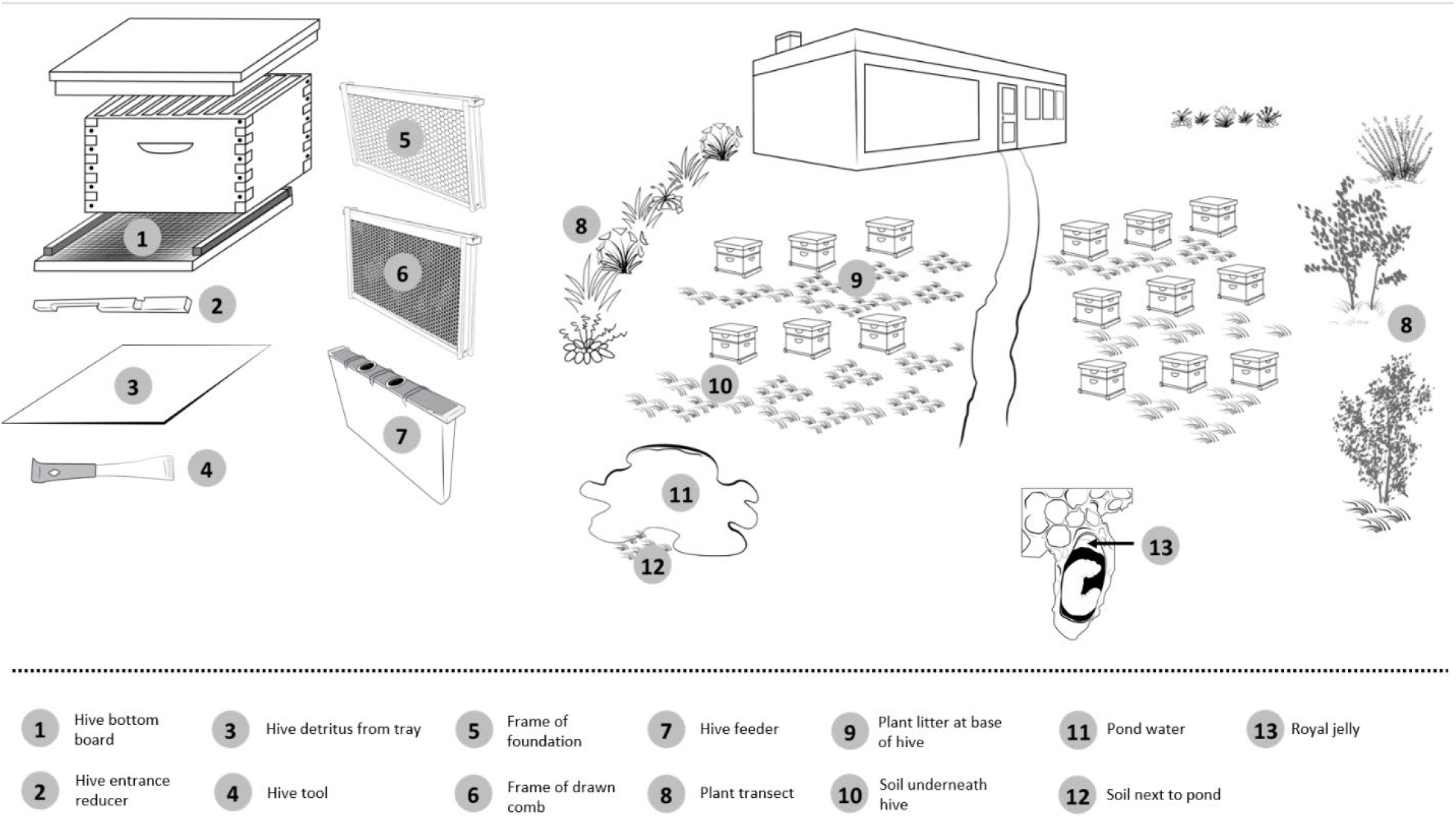
Illustration of the apiary environment from which surfaces, solids and liquids were sampled. See Table 1 for details.

#### 2.1.1 Surface samples

We obtained eDNA present on surfaces in contact with honey bees by sampling the following surfaces: hive bottom board, hive entrance reducer, hive detritus collected in corrugated plastic removable trays on hive bottom boards, hive tools, frame of foundation (Plasticell, Dadant & Sons Inc., High Springs, FL), frame of drawn comb, and plant transects (see Figure 1). All target surfaces, except drawn combs already inside the hives, were washed with 1% bleach solution and rinsed three times with activated carbon filtered water to remove all existing tissues, debris, and eDNA. Sterilized surfaces were left for 24 h to accumulate fresh eDNA before sampling occurred. The laboratory beekeepers used three new and pre-sterilized hive tools when working colonies in the apiary (n=36 hives). Each hive tool was used on one-third of the apiary (twelve colonies) before being sampled for eDNA. The beekeepers wore disposable nitrile surgical gloves to limit contamination. The 10 m plant transects, consisting of vegetation visited by honey bees close to the apiary (see Table S1 for plant species composition), were sampled after three days without rain to avoid eDNA runoff and allow accumulation of eDNA from various sources (Valentin et al., 2021).

eDNA from each surface was collected with swabs and spray/wash aggregation protocols (details follow). In addition, the frame of foundation was also sampled with a roller aggregation method (Valentin et al., 2020). Plant litter at the base of a hive was collected by directly suspending ∼ 5 g of the litter in DEPC water (#4387937, Thermo Fisher Scientific, Waltham, MA, USA) and subsequently filtering the solution. In all cases, except for the hive bottom board, swabs and spray/wash were performed on different days, or different surfaces, to obtain independent samples. The hive bottom board was swabbed first, and then the remainder of the board was sprayed/washed. Therefore, results for this surface are dependent and were combined for analyses. Frame of drawn comb samples were considered independent, despite two combs being sampled from the same hive.

Throughout all sampling, contamination was reduced by one-time use DNase/RNase free consumables, surgical gloves and collecting supplies. Field negative controls for each collection method were also taken to ensure no contamination events occurred from equipment used or from the DEPC water, as described below.

##### 2.1.1.1 Swabs

Individual sterile, DNase/RNase free forensic swabs with viscose tips (#80.634, Sarstedt, Newton, NC, USA) were used to collect surface eDNA. The swabs were moistened in UltraPure DNase/RNase free water (#10977015, Invitrogen, Grand Island, NY) to facilitate DNA transfer (van Oorschot et al., 2003) and passed over the sampling surface until residue was visible. The swabs were stored in their original sterile DNase/RNase free containers and immediately allocated into a cooler with ice packs. The swabs were refrigerated at 4°C, and DNA extraction was initiated (overnight incubation step in Proteinase K and ATL buffer) within an hour of collecting. Field negative controls consisted of a swab soaked in UltraPure distilled water prior to each sampling event.

##### 2.1.1.2 Roller

Foundation frames were sampled using a pre-sterilized (UV light + bleach 1%) sponge hand-paint roller attached to a metal handle (#144257, WHIZZ applicators 3-Piece Paint Roller Kit), dampened in DEPC water and applied across the entire frame in 3-5 successive brushes to collect eDNA. The roller was then placed directly inside a sterilized bucket, sprayed with 500 mL DEPC water to remove the eDNA on its surface, and filtered as described below [see Valentin et al., (2020) for details]. A single field negative control consisting of a sterilized foundation frame, sterilized roller and bucket, and sprayer was taken to test for contamination.

##### 2.1.1.3 Spray/wash aggregation and filtering

A new manually pressurized 5 L sprayer (X001Z2GBHT, Ivosun, Ontario, CA) was filled with 1 L of sterile DEPC water (#4387937, Santa Cruz Biotechnology Inc., Dallas, TX, USA) and operated at a consistent pressure of 36 psi (2.48 bar) to spray surfaces consistently. A 1% bleach and UV light sterilized 1.5 L collecting bucket, with an opening diameter of 13 cm, was used to collect 500 mL of spray run-off from sampling surfaces. All single use collecting equipment was replaced between samples, except the collecting bucket which was surface sterilized with a solution of 1% bleach, rinsed with filtered water three times, and UV sterilized at the end of each sampling event. Field negative controls for the sprayer and bucket were taken prior to each sample set of three replicates to ensure their sterilization and later test for contamination.

The 500 mL sample from spray/washed surfaces (i.e., the aggregate) collected in the bucket was filtered through a one-time-use eDNA filter pack composed of an outer plastic shell and an inner 0.45 µm polyethersulfone (PES) filter membrane (#11746, Smith-Root Inc., Vancouver, WA, USA). This was sufficient size to capture intracellular and intraorganellar DNA while allowing extracellular DNA and other smaller particles and organisms to pass through the filter (Turner et al., 2014). The filter pack was connected to a Pegasus Alexis field peristaltic pump (#76548, Proactive Environmental Products, Bradenton, FL, USA) via a one-time-use 1 m long Masterflex L/S^®^ Precision Pump Tubing, Platinum-Cured Silicone, L/S 15 (# EW-96410-15, Cole-Palmer Instrument Co., Vernon Hills, IL, USA) working at a speed of 52.7% (∼processing duration per sample: 5 minutes). A 500 mL sterile Nalgene bottle (Thermo Fisher Scientific, Waltham, MA, USA) used to collect the filtrate. Solid debris deposits at the base of the bucket were avoided while pumping to prevent prematurely clogging the filter. The entire filter pack (including the filter containing embedded sampled eDNA) was aseptically stored in a cooler with dry ice packs, using single use surgical gloves, for <1 hour before DNA extraction was initiated (overnight incubation step in Proteinase K and ATL buffer at 65°C). The 500 mL filtrate containing material smaller than 0.45 µm (including viruses) was immediately placed in a cooler with ice, and then stored at -80°C for future research.

Samples from drawn comb contained runoff honey when spraying DEPC water into the comb. Two filters were used per drawn comb as the honey completely saturated filters prior to filtering 500 mL of the collected spray runoff. These two filters were then combined for extraction. In addition, plant litter at the base of the hives was collected (*circa* 5 g) and 500 mL of DEPC water was added to suspend any surface eDNA present into solution. After 1 min, the water was filtered as described previously, avoiding solid debris to prevent clogging the filter. As with the other filtering methods, a sterilized bucket and an unopened filter pack were used to collect a field negative control prior to sample collection as previously described.

#### 2.1.2 Physical sample collection

Physical samples were directly collected from around the apiary. We obtained suspended eDNA in water that honey bees contacted by sampling 500 mL of fructose:water (1:1) from in-hive feeders (Mann Lake Ltd., Hackensack, MN) and 500 mL of water from water sources in the vicinity of the apiary that honey bees often visited for water collection (i.e., a permanent pond of 1,885.7 m^2^; a decorative pond of 30.24 m^2^ ; and a temporary retention pond of 496.36 m^2^). The liquid samples were filtered as described above, although filtration took longer (∼20 minutes) for in-hive feeders due to fructose water’s high viscosity. Solid soil samples (∼5 g) were collected from soil underneath the hive and soil near the edge of the pond using a one-time use sterile spatula. The samples were placed in 50 mL DNase/RNase free tubes (# 62.547.254, Sarstedt, Newton, NC, USA) which were placed in a cooler with ice until DNA extraction. The sampled royal jelly was purchased from a commercially available US supplier that imported it from the Zhejaing Region of China. The 50 g of royal jelly was equitably split into four 50 mL tubes.

### 2.2 DNA extraction and quantification

#### 2.2.1 Swabs and filters

An individual viscose tip (0.5 × 1 mm) swab was cut at the edge of its polystyrene stick using flame sterilized scissors (i.e., Bunsen Burner for 10 s followed by 95% ethanol dip and Bunsen again for 2 s), and then placed in a 2 mL sterile DNase/RNase Biosphere® plus free screw-cap tube (#72.694.217, Sarstedt, Newton, NC, USA). The entire filter (4.7 cm Ø) was removed from the filter pack outer shell under a hood, placed in a sterile petri-dish with sterile forceps, and shredded with a one-time use sterile blade. Sequential vertical stripes of 3 mm were made across the entire filter. The shredded filter was then transferred into a 2 mL sterile DNase/RNase Biosphere® plus free screw-cap tubes (Sarstedt, Newton, NC, USA). DNA was extracted using a DNeasy® Blood and tissue kit (#69504, Qiagen, Valencia, CA, USA), with a slightly modified version of the Renshaw et al. (2015) protocol for filter membranes plus extraction solution to fit into the 2mL tubes.

Borrowing from Renshaw et al. (2015), swabs and filters in 2 mL screw-cap tubes containing one 5 mm Ø stainless steel bead (Qiagen, Valencia, CA, USA) were completely immersed in 63 μL of Proteinase K and 567 μL of buffer ATL. The samples were homogenized at 4 M/s for two sequential steps of 5 s (Benchtop FastPrep-24™ Homogenizer (MP Biomedicals LLC, Santa Ana, CA, USA)). The tubes were briefly centrifuged for 5 s and incubated overnight in a rocking water bath at 65 °C. After incubation, 500 μL buffer AL and 500 μL 100% ethanol were added to the tube. The lysate was then passed through the spin columns (three iterations were required to pass all the lysate through a column), and the subsequent extraction steps followed the manufacturer’s recommendations. Total DNA was eluted in a volume of 40 μL and stored at -20 C. All pipette tips were sterile and DNase/RNase free. The handling tools were replaced between consecutive samples.

#### 2.2.2 Soil

Soil samples were processed using OMNI Soil Purification Midi kit (26-016, PerkinElmer inc., Waltham, MA) following the manufacturer’s protocol and overnight incubation at 65°C after adding SD buffer, to enhance lysis. Total DNA was eluted in a volume of 40 μL.

#### 2.2.3 Royal jelly

DNA was extracted from royal jelly in a USDA Level II Biological Containment Facility (USDA-PPQ permit # P526P-21-01762) using a modified protocol by Bovo et al. (2020). Briefly, 40 mL of ultrapure water (Invitrogen, Grand Island, NY) was added to the tubes containing royal jelly. The tubes were vortexed for 1 min and incubated for 10 min at 65° C in a water bath. Royal jelly monosaccharide sugars were removed by centrifugation of the tubes at 4,000 rpm for 25 min (Centrifuge 5810 R, Eppendorf AG, Hamburg, Germany). The supernatant was discarded and the pellets from the four tubes were merged into a 50 mL tube that was centrifuged for an additional 25 min. The additional supernatant was removed and 700 µL of the pellet was transferred to each of five 1.5 mL Eppendorf^®^ tubes pre-filled with 250 µL glass beads (#19-642-3, OMNI International Inc., Kennesaw, GA). The tubes were vortexed for 3 min and then centrifuged for 1 min at 12 000 rpm (Sorvall Legend Micro 21R, Thermo Fisher Scientific, Waltham, MA). Then, 700 µL of pellet, without the beads, from each tube was transferred into a new Eppendorf tube. 500 µL of ATL buffer (Qiagen, Valencia, CA) and 60 µL of Proteinase K (Promega, Madison, WI) were added to each tube, vortexed for 1 min, and incubated overnight in a water bath at 60°C. A no template control (NTC) using ultra-pure water was added at this stage.

DNA was extracted from royal jelly lysed products, as well as the NTC, using a DNeasy® Blood and tissue kit (Qiagen, Valencia, CA, USA). Each incubated royal jelly product was divided into two new Eppendorf tubes. A 1:1 mix of 800 µL 100% ETOH:Buffer AL was added to each of the tubes. The content of each pair of tubes was filtered through a single DNeasy® membrane, washed with 500 µL buffer AW1 and 500 µL buffer AW2, respectively, and eluted in 40 µL 1:1 mix of Elution buffer + ultrapure water. After quantification, the royal jelly DNA was pooled and speed vacuumed using a Vacufuge Plus (Eppendorf, Hamburg, Germany) for 30 mins to concentrate DNA.

#### 2.2.4 Positive DNA yield control

Fresh, randomly collected *A. mellifera* specimens from colonies at the HBREL were extracted for use as PCR positive controls. The bees were dissected with a vertical cut along the middle section of the entire thorax and head to facilitate lysis. The tissues were submerged in 10 µL Proteinase K and 180 µL Buffer ATL, in 2 µL screw-cap tubes containing one 5 mm Ø stainless steel bead. The samples were homogenized at 4 m/s for two sequential steps of 5 s (Benchtop FastPrep-24™ Homogenizer (MP Biomedicals LLC, Santa Ana, CA, USA). The tubes were briefly centrifuged for 5 s and incubated overnight in a rocking water bath at 65 °C. The extractions followed the Qiagen’s DNeasy® Blood and tissue kit manufacturer’s instructions for tissues.

#### 2.2.5 DNA quantification

DNA quantification was determined using a Qubit™ dsDNA HS Assay Kit (#Q33230, Thermo Fisher Scientific, Waltham, MA, USA) with an Invitrogen™ Qubit™ 3.0 Quantitation Fluorometer, following manufacturer’s recommendations. Total DNA was stored at -20°C prior to PCR and sequencing.

### 2.3 Metabarcoding

#### 2.3.1 Primers

We enriched invertebrate 16S and Cytochrome Oxidase I (COI) mitochondrial DNA (mtDNA) using the IN16STK primer set (da Silva et al., 2019) and FwCOI primer set fwhF2 and fwR2n (Vamos et al., 2017) respectively. Microbial primers for bacterial V4 16S (Gohl et al., 2016) and fungal ITS (Olimi et al., 2022) metabarcoding were also used to assess the non-viral microorganisms present within apiaries. All primer sets were indexed with 8 bp tags that would result in unique dual index combinations per reaction per primer set, following Taberlet et al. (2018), and available as Table S2.

#### 2.3.2 PCR

The samples were enriched in triplicate for each primer set, resulting in twelve replicates per sample across four libraries, all with unique dual index combinations (See Supplementary File 1 for primer plate plans). Each 96-well plate included the entire set of samples (Table S2) and each individual well contained 20 µL PCR reactions with a PCR bead (#27955701, Cytiva illustra™ PuReTaq Ready-To-Go™ PCR Beads), 9 µL of UltraPure™ DNase/RNase-Free Distilled Water, 4 µL of each forward and reverse indexed primer at 1 mM concentration, and 3 µL of sample template. We opted to use PCR beads as attempts with different polymerases and mixes in the marker (Hot Start AmpliTaq Gold DNA; Thermo Fisher Platinum II HotStart PCR MasterMix; OneTaq Hot Start Master Mix 2X) would not amplify across all sample types. The PCRs plates were run on an Eppendorf 6331 Mastercycler® as follows: 95°C for 2 min (initialization), followed by 40 cycles of 95°C for 30 s (denaturation), 52°C for 30 s (annealing), and 72°C for 50 s (extension). A final elongation step at 72°C for 2 min was used. Samples were stored at 4°C prior to gel electrophoresis in 1% agarose gels with 1 µL SYBR Safe dye (# S33102, Thermo Fisher Scientific, Carlsbad, CA) for 35 min at 150 V. The resulting gels were visualized using a BIO-RAD ChemiDoc™ XRS+ imaging system (Hercules, CA, USA).

After confirming successful PCR enrichment from a subset of samples, all reactions from each plate were evenly pooled into DNA-free 1.5ml microcentrifuge tubes (i.e., 12 tubes in total, three per primer set). All pools were purified using a Qiagen MinElute PCR Purification Kit using the manufacturer’s recommended protocol, then combined by primer set into four final superpools (for Arthropod COI; Arthropod 16S, Fungi ITS, and Bacteria 16S, respectively). Each of these four superpools were submitted to the University of Florida Interdisciplinary Center for Biotechnology Research Genomics Core (UF-ICBR) for final size selection to remove any non-specific amplifications and complete library preparation before being sequenced on an Illumina MiSeq™ using the Illumina v3 600-cycle reagent kit for paired-end sequencing.

### 2.4 Bioinformatics

#### 2.4.1 Arthropod Bioinformatics – OBITools

All arthropod bioinformatics were conducted using the OBITools software package (Boyer et al., 2016) by first aligning paired end reads, then assigning merged sequences back to their original PCR reactions via the tagged primer’s unique dual index. No errors were allowed when matching the primer tags and no more than two errors when matching the primer sequences. Sequences with ambiguous bases, those that were not merged, and those outside the expected size range (i.e., 70–220 bp) were filtered out of the dataset. Sequences were then dereplicated and read singletons were filtered out from the dataset. Taxonomic assignments were made using a global reference database made from EMBL (release 143). To account for potentially erroneous sequences generated either through PCR enrichment or sequencing, the *obiclean* function was used (parameters *d*=1 and *r*=0.25) to assign all sequences as either being a true sequence (‘*head*’), variant of a true sequence (‘*internal*’), or not associated with any other sequence (‘*singleton*’). A molecular Operational Taxonomic Unit (mOTU) × PCR reaction table was then outputted for further quality filtering.

Further filtering of the dataset was performed in R (v 4.1.3 - (R Core Team, 2013)) by first removing mOTUs with low read depth based on the greater of two thresholds: *(i)* <10% the average read depth per PCR reaction, and *(ii)* at least 1,000 reads per reaction. mOTUs with poor taxonomic assignment scores <80% (indicative of chimeras) were also filtered out of the dataset (Pansu et al., 2015). Sequences were identified as being contaminants if their average relative max frequency was within any of the various controls throughout the study and discarded from the dataset. PCR replicates with high levels of contaminant sequences and high dissimilarity among replicates from the same sample were also discarded. The sample was removed from the dataset if no more than one PCR replicate per sample remained post filtering. Once our filtering steps were complete, PCR replicates were aggregated to sample (i.e., island transects) by average number of reads. As a final measure, any mOTUs in the corresponding data table with <1% of the respective reads within the overall sample were discarded.

#### 2.4.2 Microbial Bioinformatics – DADA2

All microbial bioinformatics were performed using the DADA2 pipeline version 8 (Callahan et al., 2016). Forward and reverse fastq files were demultiplexed and renamed into unique forward and reverse FASTQ files per reaction name using barcode and pattern file made from cutadapt (Martin, 2011). All FASTQ files (grouped by either forward or reverse read) were assessed for average read quality throughout the length of the sequence, with the length of the read trimmed to remain above a quality score of 30 using the *filterANDtrim* command in DADA2 to remove primers and low-quality portions. While this is typically not advised for ITS sequences due to the variable length of the locus (Collins & Paskewitz, 1996; Hill et al., 2008), the fungal library was size selected to yield a consistent amplicon length. Filtered files had a maximum of two expected errors and no Ns allowed. Forward and reverse read files were then merged, dereplicated, and an Amplicon Sequence Variant (ASV) × PCR reaction table generated for further quality filtering. Chimeras were removed from the ASV × PCR reaction table, and were then used to assign and output taxonomic assignments for each ASV within the dataset using the Silva v138 (Yilmaz et al., 2014) reference database. Species taxonomic assignment were made using the Silva species assignment v138.

Further processing and filtering of the dataset was performed in R by first merging both DADA2 file outputs into a phyloseq object using the phyloseq package (v 1.38.0 – (McMurdie & Holmes, 2013)). Contaminant sequences were filtered out using the decontam package (v. 1.14.0 – (Davis et al., 2018)), and ASVs not identified within the kingdom bacteria as observed within the Silva taxonomy were removed. PCR replicates with high dissimilarity were also discarded and PCR reactions with a read depth below 1000 reads were removed. If no more than one PCR replicate per sample remained post filtering, the sample was removed from the dataset. Once complete, replicates were merged by sum and the entire dataset rarefied to an even read depth. Samples were rarified to 547 reads for arthropod COI, 909 reads for arthropod 16S, 4341 for bacteria 16S, and 2744 for fungi ITS.

### 2.5 Analysis and visualization

The arthropod datasets (i.e., arthropod 16S and COI) were each rarefied to the respective sample with the lowest read depth before proceeding with calculations. Because of the interdependency of the hive bottom board samples (HBB, Table 1) being taken from the same hive at the same time, these samples were aggregated by sum across all four datasets prior to performing any analysis. mOTU and ASV richness was calculated across all four datasets (i.e., arthropod 16S and COI, bacterial 16S, and fungal ITS) for the following categories: surface sampled (Table 1), the type of sample that was taken (e.g., Table S1), and the source of the sample (i.e., hives, transects, or soil). We calculated within sample dispersion via Bray-Curtis dissimilarity and assessed whether pairwise differences in dissimilarity were significant via analysis of variance (ANOVA). Community composition for each of the four datasets were visually assessed via Non-metric Multi-Dimensional Scaling (NMDS), with groupings based on sample type, collection type, and a combination of the two. Visual assessment of the sample + collection ordinations allowed us to select for five likely informative combinations. A binary factor was added to the metadata indicating whether or not these five selected sample type + collection methods were selected (1) or not selected (0) to test via PERMANOVA whether they significantly explain the variation in the data. Composition of surface samples, collection methods, and source of samples for taxonomic class, order, family, and genus were also visually assessed across all four datasets via stacked bar plots, taking the top 42 mOTUs for arthropod 16S (explanation in results) and the top 30 mOTUs for the remaining three datasets.

## 3. Results

### 3.1 Sampling locations

Most of the thirteen surfaces, liquids, and solids sampled from the apiary environments generated viable eDNA for each primer set. The main exception being soil next to pond (no data obtained). We were also unable to obtain results for either arthropod primer set for soil underneath hive; and hive feeder failed to provide data for arthropod 16S and fungi.

The richness plots presented in the supplementary materials for each primer set are stacked, summing the data collected with different methods for each sample type. Greater richness does not necessarily mean more taxonomic information was collected as multiple variants for the same taxonomic assignation or variants occur; yet, these provide some indication of the data (Figure S1a, S1b, S2a, S2b). Where more than one collection method was used for a sample type, they facilitate a visual comparison of richness obtained with each method. Plant transects, hive bottom board, entrance reducer, and pond water had the highest absolute richness for arthropod COI data (>25; Figure S1a), while hive bottom board and plant transects were the highest for arthropod 16S (38 and 24 respectively; Figure S1b). For bacteria 16S, the pond water had the highest absolute richness (>450), followed by hive tools, hive bottom board, and frame of foundation (>300; Figure S2a). For fungi ITS, several sample types had high absolute richness (>125) including entrance reducer, hive tools, plant litter, pond water and hive bottom board (Figure S2b). In all four datasets, royal jelly had the lowest absolute richness.

The dispersion results by sample type show how similar replicates were, with a lower score indicating higher repeatability. For arthropod COI, replicates of hive detritus from tray were the most similar, while replicates from plant transects varied most (Figure S1e). For arthropod 16S, hive detritus from tray, hive entrance reducer and frame of drawn comb were the most repeatable, while plant transects were again least similar (Figure S1f). For bacteria, frame of drawn comb was the most repeatable, followed by hive feeder and plant litter at base of hive (Figure S2e). Frame of drawn comb was again the most repeatable with fungi ITS (Figure S2f), while for both bacteria and fungi, pond water was least similar. Our pairwise assessment of dispersion among samples yielded some significant differences across all four datasets (Table S3).

### 3.2 Sampling methods

We collected eDNA using both swabs and spray/wash methods for hive bottom board, hive entrance reducer, hive detritus, hive tools, frame of foundation, frame of drawn comb, and plant transects. In addition, we also collected eDNA using a roller for frame of foundation. This allows some comparison of these methods for these sample types. For arthropod COI and 16S data, bacteria 16S, and fungi ITS datasets, filters from spray/wash method had more absolute richness than did swabs (Figure S1a, S1b, S2a, S2b). The composition of relative read abundances also differed dramatically between collection types (Figure 2a, 3a, 4a, 5a), with spray/wash being more diverse. The NMDS ordination plots show that swabs and spray/wash filters collected overlapping DNA for each primer set, this indicated by overlapping ellipses (Figure S1c, S1d, S2c, S2d).

**Figure 2.**
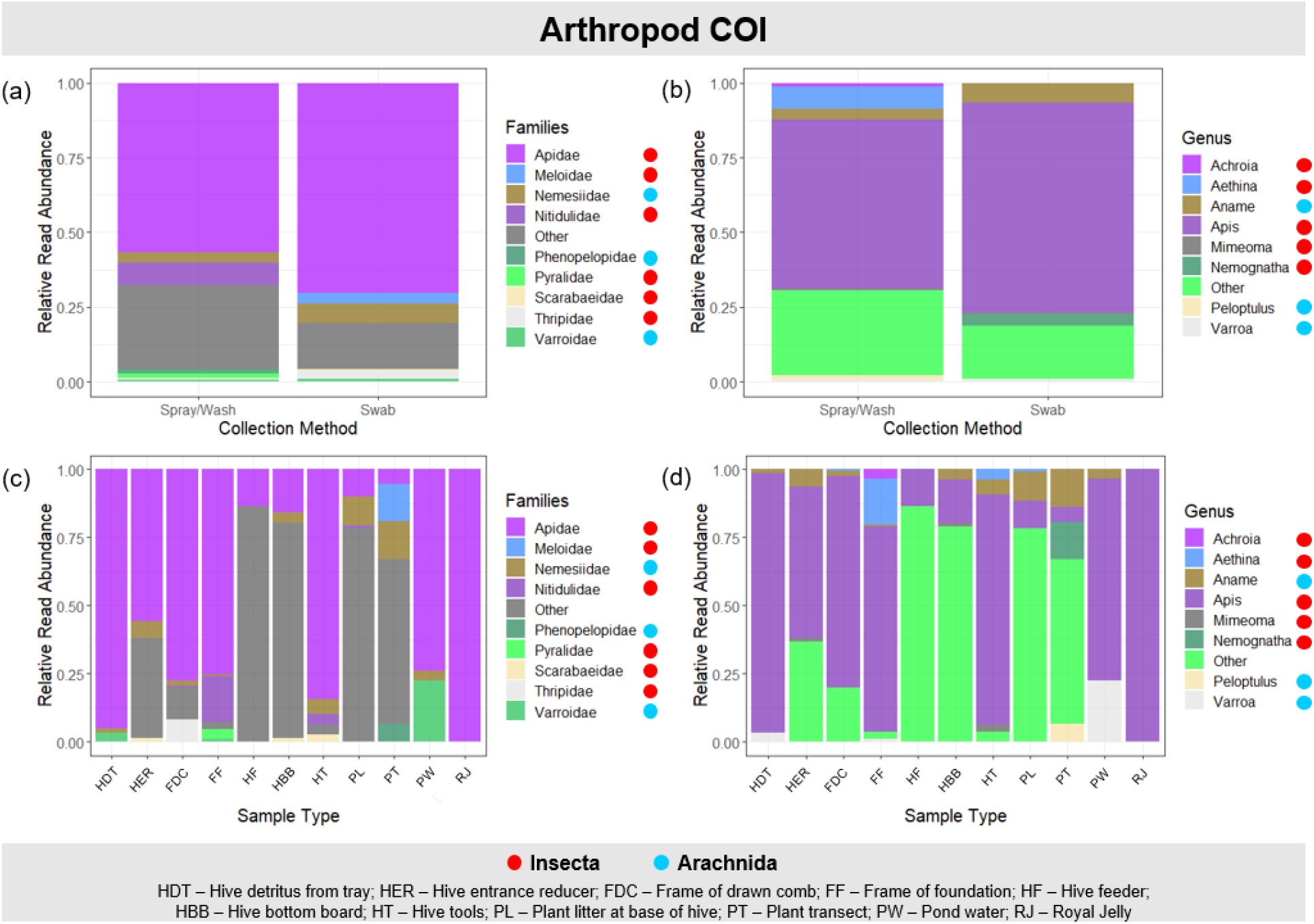
Results of the top 30 molecular Operational Taxonomic Units (mOTU) representing 83.94% of the mOTUs for the arthropod COI region. Relative Read Abundances of mOTUs are shown at the Family (a, c) and Genus (b, d) levels and are separated per collection method (a, b) and sample type (c, d). Samples not shown were discarded during the bioinformatic pipeline.

**Figure 3.**
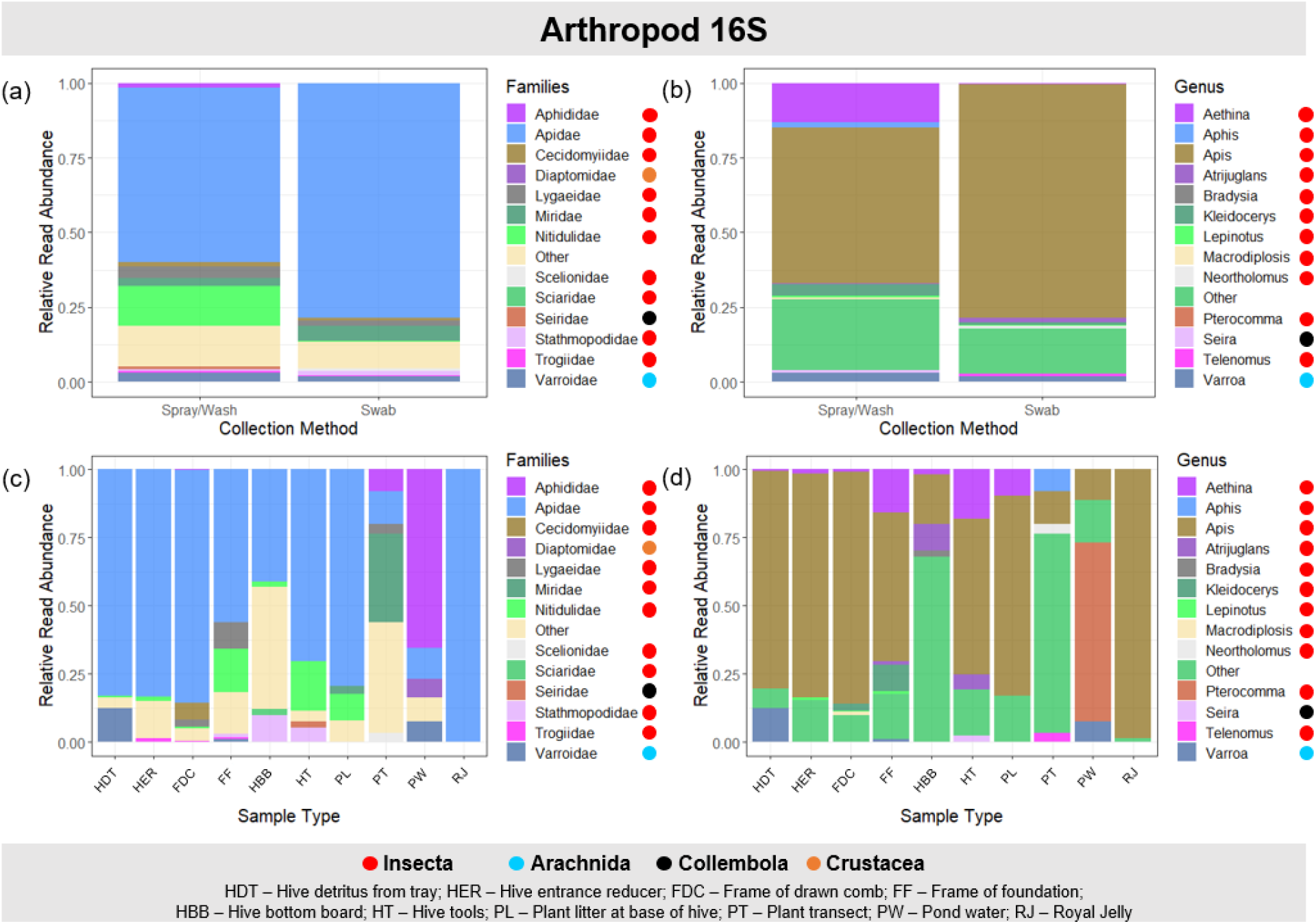
Results of the top 42 molecular Operational Taxonomic Units (mOTU) representing 96.47% of the mOTUs for the arthropod 16S region. Relative Read Abundances of mOTUs are shown at the Family (a, c) and Genus (b, d) levels and are separated per collection method (a, b) and sample type (c, d). Samples not shown were discarded during the bioinformatic pipeline.

**Figure 4.**
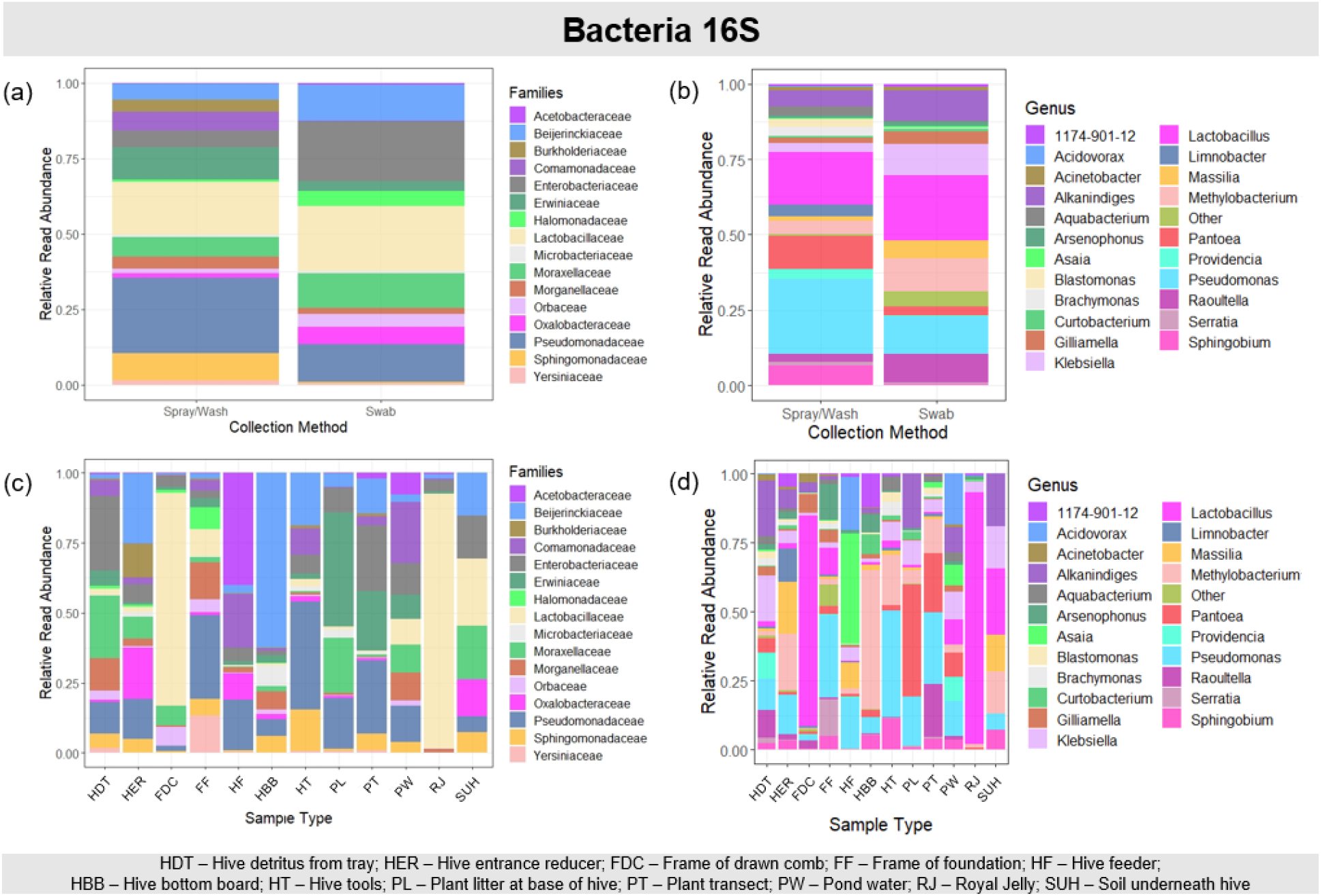
Results of the top 30 amplicon sequence variants (ASV) representing 54.87% of the data for the bacterial 16S region. Relative Read Abundances of ASVs are shown at the Family (a, c) and Genus (b, d) levels and are separated per collection method (a, b) and sample type (c, d). Samples not shown were discarded during the bioinformatic pipeline.

**Figure 5.**
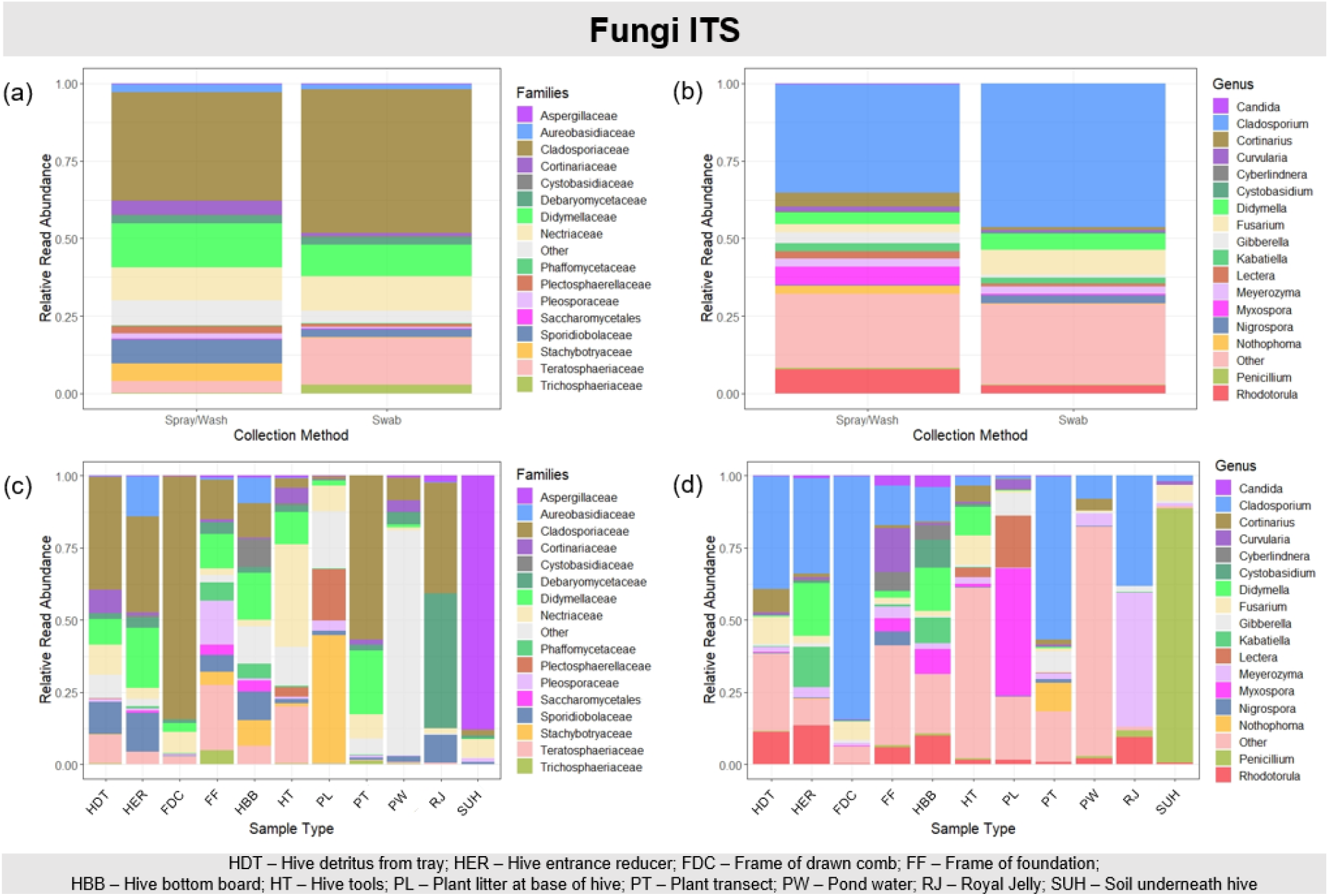
Results of the top 30 amplicon sequence variants (ASV) representing 67.17% of the data for the fungal ITS region. Relative Read Abundances of ASVs are shown at the Family (a, c) and Genus (b, d) levels and are separated per collection method (a, b) and sample type (c, d). Samples not shown were discarded during the bioinformatic pipeline.

### 3.3 Species detected based on eDNA metabarcoding

#### 3.3.1 Arthropods

In both datasets, *Apis* spp.-related molecular Operational Taxonomic Units (mOTUs) were abundant. We have chosen to present data with *Apis* spp. results as this paper is exploratory, and in some cases *Apis* spp. may be the target of eDNA metabarcoding (see discussion). Alternative results with *Apis* spp. excluded are available in the supplementary materials (Figure S3, S4).

##### 3.3.1.1 COI DNA region

The top 30 mOTUs representing 83.94% of the data are shown in Figure 2. As expected, the family Apidae, and genus *Apis*, dominates all collection types. Different samples collected different targets (Figure 2c, 2d). Notably, DNA from *V. destructor* was detected in samples from board under hive and pond water (Figure 2c), while evidence for small hive beetles was found in frame of foundation and hive tool samples (Figure 2d).

The total filtered mOTUs for arthropod COI were 150 (Table S4). Amongst these, we found DNA evidence of honey bees *Apis* sp. (n=1), *Apis mellifera* (n=1); small hive beetles (n=2); *V. destructor* sp (n=1); and greater (n = 1) and lesser (n = 1) wax moths. We also detected a mOTU from *Brachymyrmex* sp. of rover ants (n=1), and *Aname* sp. MYG271 (n=17) an endemic Australian spider genus. Detecting DNA does not mean that this Australian spider is present in the UF HBREL apiary. As over 100 sequences are currently unclassified *Aname* sp. on NCBI Taxonomy Browser (NCBI:txid2625272), it may be a closely related US species that does not have COI data available (see discussions in Beng & Corlett, 2020).

##### 3.3.1.2 16S DNA region

The top 42 mOTUs representing 96.47% of the data are shown in Figure 3. We chose to increase the number of mOTUs displayed, as *Apis* data took up 12 mOTUs itself. Similar to COI data, *Apis* data were predominant, and *V. destructor* DNA was detected in samples from board under hive and pond water (Figure 3c, 3d). DNA from small hive beetles was found in frame of foundation and hive tools (like COI), as well as in samples of plant litter at base of hive (Figure 3d).

The total filtered mOTUs for arthropod 16S were 91 (Table S4). Similar to COI dataset, we found *Apis mellifera* (n=3), small hive beetle (n=4), *V. destructor* (n=2), greater wax moth (n=1), and *Brachymyrmex* sp. of rover ants (n=2). Unlike with the COI primer set, we did not detect the lesser wax moth, nor the *Aname* sp. of spider, with this primer set.

#### 3.3.2 Bacteria 16S DNA region

The top 30 ASVs representing 54.87% of the data are shown in Figure 4. The total filtered ASVs for Bacteria 16S were 2437. We detected DNA from bacterial species of apiculture interest. These were: a) European foulbrood bacterium *Melissococcus plutonius* (n=1), commonly reported in hives by visual observation; b) *Bombella apis* (n=1), previously reported in the midgut of honey bees as a possible symbiont (Yun et al., 2017); c) *Bartonella apis* (n=1), a midgut symbiont (Kešnerová et al., 2016) and d) *Wolbachia* sp. (n=2). *Wolbachia* sp. is an intracellular parasite in many arthropods (Hilgenboecker et al., 2008; Jeyaprakash & Hoy, 2000) that preferentially infects testes and ovaries, with implications in fecundity (Werren, 1997), and commonly found in *V. destructor* infesting hives (Grau et al., 2017).

We did not detect American foulbrood (*Paenibacillus larvae*), but we detected DNA from five *Paenibacillus* spp. in the dataset. Only one of these could be resolved to species level and was identified as *P. gelatinilyticus*. Thus, our method would most probably have detected this serious hive disease if it were present in the samples.

#### 3.3.3 Fungi ITS DNA region

The top 30 ASVs representing 67.17% of the data are shown in Figure 5. The total filtered ASVs for Fungi ITS were 804. DNA from mycotoxin-producing fungi genera that are associated with bees (e.g., *Aspergillus, Penicillium*, and *Fusarium* (Keller et al., 2014)) was detected. We detected DNA from *Aspergillus conicus* Blochwitz 1914 (n=1); *A. insuetus* (Bainier) Thom & Church 1929 (n=1); *A. flavipes* (Bainier & Sartory) Thom & Church 1926 (n=1); *A. flavus* Link 1809 (n=1); *Aspergillus* spp. (n=2); *Penicillium citrinum* Thom 1910 (n=1); *P. cinerascens* (Piper) Rydberg 1908 (n=1); *P. brevicompactum* Dierckx 1901 (n=1); *P. polonicum* Zalessky 1927 (n=1); *Penicillium* sp. (n=1); *Fusarium acutatum* Nirenberg & O’Donnell 1998 (n=3); *F. longipes* Wollenw & Reinking 1925 (n=1); *F. chlamydosporum* Wollenw & Reinking 1925 (n=2); *F. polyphialidicum* Malasas 1986 (n=1); *F. solani* (Martius) Sacccardo 1881 (n=1); *F. kyushuense* Aoki & O’Donnell 1998 (n=1); *F. sporotrichioides* (Sherb.) 1915 (n=1); and *Fusarium* sp. (n=3). We did not detect DNA from chalkbrood disease causing *Ascosphaera apis*, nor *Bettsia alvei* (beehive mold on stored pollen). We also did not detect DNA from *Nosema ceranae*, a gut pathogen of honey bees (additional discussion below regarding *N. ceranae*).

### 3.4 Most informative sample type and collection method

We combined the data from a sample type + collection method to determine which sample type and collection method were able to explain most of the data for each primer set. The five selected sample type + collection methods for COI were hive tools (swab), plant litter at base of hive (spray/wash filter), hive entrance reducer (spray/wash filter), hive bottom board (combined methods), and frame of foundation (spray/wash filter). They significantly explained the variation in the data (PERMANOVA, P = 0.028), accounting for 50.90% of the mOTUs in the dataset, which in turn covered 53.33% of all identified orders, 37.04% of identified families, and 42.31% of all genera. For arthropod 16S, the selected samples that significantly explained the variation in the data (PERMANOVA, P = 0.013) were spray/wash filters from frame of drawn comb, hive tools, and frame of foundation, together with swabs from frame of drawn comb and hive bottom board (combined methods). They covered 60.61% of the mOTUs in the dataset, which in turn covered 80% of all identified orders, 69.57% of identified families, and 69.57% of all genera.

For bacteria, swabs from hive detritus from tray, frame of foundation, hive entrance reducer, and hive tools, together with hive bottom board (combined methods). Collectively, these five samples covered 41.44% of the ASVs in the dataset, which in turn covered 61.49% of all identified orders, 57.61% of identified families, and 52.88% of all genera (PERMANOVA, P = 0.002). The five selected sample type + collection methods that significantly explained the variation in the fungal data (P = 0.01) were frame of foundation (spray/wash filter), hive entrance reducer (spray/wash filter), frame of drawn comb (swab), frame of drawn comb (spray/wash filter) and hive bottom board (combined method). These covered 35.45% of the ASVs in the dataset, which in turn covered 70.37% of all identified orders, 64.95% of identified families, and 54.67% of all genera.

## 4. Discussion

In most cases, the different surfaces yielded some DNA, except for some soil samples (likely due to the lack of availability of our first-choice extraction kit). The results of the swab and spray/wash technique were similar when they could be applied to the same surface, and all four primer sets performed well. We were able to detect DNA from *A. tumida, V. destructor* and European foulbrood, all of which matched our visual observations of clinical signs of these pests and pathogens in the hives we tested.

Naturally, DNA from some species was only found in specific locations, which is expected based on the locations and behaviors of different organisms. Collectively, our data show that eDNA metabarcoding is a viable method for monitoring honey bee pests and pathogens.

### 4.1 Suggestions for sampling locations and methods

Different organisms were detected in different samples, which is important to keep in mind when developing future methods using eDNA to detect honey bee pests and pathogens. Therefore, the choice of sample type and collection method may be dependent on the organism targeted using eDNA monitoring. Logically, surfaces that were ranked as easy to sample (Table S1) would be best to sample for large-scale monitoring. For example, requiring hives be lifted to sample hive bottom boards is not feasible at larger scales, and would require too much additional time from beekeepers. The easy-to-sample surfaces included: hive entrance reducer, hive detritus from tray, plant litter at base of hive, and water from nearby pond that honey bees visit. Hive tools were an excellent harborage of eDNA and provided useful information on arthropod and microbial communities. Sampling hive tools after use in routine beekeeping management may help alleviate the otherwise high sampling effort in this study (requiring new tools, gloves and working multiple hives).

Spray/wash filters worked marginally better than did swabs (in terms of higher absolute richness). Our PERMANOVA results selected for spray/wash filter samples when targeting arthropod DNA, while swabs performed best when targeting bacterial DNA. Importantly, most organisms were detected by both methods when tested on the same surfaces. Given the ease of eDNA collection with swabs and the affordable cost associated with using them, they may provide the best option for further development of this method.

Unfortunately, we were unable to evaluate the information available in soil underneath hive and soil next to pond fully. Most DNA from soil extraction did not make it into final dataset as the read depths were too low. It is possible that the quality of the DNA extraction was not optimal in our study. COVID-19 constrained the availability of our first-choice extraction kits. Future research could expand on this, especially as eDNA collected from plant litter at base of hive was informative.

### 4.2 Discussion of species found

Our data matched the visual hive observations for small hive beetles, *V. destructor* and European foulbrood. No wax moths were seen, but wax moths are cryptically present in many hives (Sohail et al., 2017), likely including the ones we sampled. We were able to detect DNA from European foulbrood, which suggests that if American foulbrood were present in the test colonies, our methods would be able to detect it. As expected, our data suggest that honey bees interact with several other organisms in their hives and immediate environment.

Two interesting organisms from which we detected DNA were *Brachymyrmex* sp. and *Wolbachia. Brachymyrmex* sp., or rover ants, DNA was found using arthropod COI and 16S primers. These ants have been found to carry pathogenic viruses of bees (e.g., DWV; BQCV; SBV) (Payne et al., 2020). *Wolbachia* is an intracellular parasite in many arthropods (Jeyaprakash & Hoy, 2000), preferentially infecting testes and ovaries with implications in fecundity (Werren, 1997). It has been found in *A.m. capensis* (Eschsholtz 1822) eggs but not *A.m. scutellata* (Lepeletier 1836). Potential involvement of *Wolbachia* sp. on *A. m. capensis* thelytoky needs to be explored further (Jeyaprakash et al., 2009).

### 4.3 Unobserved or “missing” target species

We did not detect *Nosema* spp. intracellular parasites in any of our samples, as the metabarcoding primers utilized do not amplify *Nosema* spp. in silico. This is not surprising as *Nosema* (Microsporidia: Nosematidae) was formerly considered a Protozoa with no mitochondria (Germot et al., 1997), hence, closely related to but outside the Kingdom Fungi (James et al., 2006; Y. Liu et al., 2009; Y. J. Liu et al., 2006). However, the taxonomic position of this phylum is complex and its members (including *N. ceranae* – (Cornman et al., 2009) are commonly reported as Fungi (Hibbett et al., 2007)).

To determine if *Nosema* DNA was collected, and could be targeted by eDNA barcoding, we ran a basic molecular detection PCR (Fries et al., 2013) using Nos-16S primers (Stevanovic et al., 2011) to amplify a 488bp fragment of mitochondrial 16S. These specific *Nosema* spp. primers are used in the APHIS National Honey Bee Disease Surveys. Using leftover eDNA samples from the hive bottom board (swab and filter, samples 71 and 73 in Table S1), a PCR protocol was performed as described in our methods section 2.3.2 using individual tubes with a bead (#27955901, Cytiva illustra™ PuReTaq Ready-To-Go™ PCR Beads), instead of a 96 plate. Amplicons were identified using BLAST (Altschul, 1997) as *N. ceranae*, and submitted to GenBank (Accession numbers: OP325565, OP325566). Therefore, future eDNA studies could target *Nosema* spp. using targeted primers on the collected eDNA samples.

There are at least 14 arthropods known to be problematic for honey bees (Boncristiani et al., 2021). *Braula* spp. and the parasitoid *Rondaniooestrus apivorus* Villanueve 1916 would not be detected by the arthropod FwhCOI primers used. These species have no reference sequences, which is a common problem with eDNA metabarcoding (see discussions in (Beng & Corlett, 2020)). All other common arthropod pests of honey bees are able to be detected by this primer set. The IN16STK2 primer set would not detect *Acarapis woodi* (tracheal mite), *Apocephalus borealis, A. grisella, Braula* spp., *Megaselia rufipes, R. apivorus, Senotainia tricuspis*, and *Tropilaelaps* spp. This highlights an additional need to bolster the reference database for these honey bee-related species to account for missing targets.

### 4.4 Royal jelly

Royal jelly is known to have antimicrobial properties against microorganisms by the activity of its peptides, enzymes, and fatty acid, such as Royalisin, 10-hydroxy-2-decenoic acid, Jelleines, and Major Royal Jelly Proteins (Fratini et al., 2016; Fujiwara et al., 1990; Harwood et al., 2021). Its antimicrobial activity also includes certain species of molds and fungi (Fujiwara et al., 1990). The previously reported microbiome in royal jelly includes the bacteria *Acetobacteraceae Alpha 2.2, Lactobacillus kunkeei, Stenotrophomonas* spp., *Rhodanobacter* spp., as well as bacteria of the bee gut microbiome (e.g., *Xanthomonadaceae, Actinobacteria, Enterobacteriaceae* and *Lactobacillus* spp.) (Kafantaris et al., 2021). The results of our study did confirm DNA from some of those bacteria; DNA from four gram positive *Lactobacillus* spp. bacteria were found. Lactobacilli are considered to be common beneficial bacteria of the honey bee gastrointestinal tract (Nowak et al., 2021) and have antifungal properties (Karami et al., 2017). Some other bacterial findings include DNA from *Gilliamella apicola*, which has the ability to ferment sugar and inhabits areas in the honey bee directed toward the center of the lumen (Nowak et al., 2021), and gram negative *Bombella apis*, that inhabits the honey bee mid gut (Nowak et al., 2021). Interestingly enough, DNA from *M. plutonius* (European foulbrood), a honey bee pathogen that causes a severe larvae enteric disease, was also found (Forsgren, 2010; Ory et al., 2022). DNA from one fungi, *Penicillium citrinum*, known to have antimicrobial activity, was also found (Mazumder et al., 2002). Other fungal DNA found included *Aspergillus sp*., an ubiquitous and known mycotoxin contaminant of food and produce (Navale et al., 2021); *Fusarium kyushuense, which* produces aflatoxin, a vigorous mycotoxins (Schmidt-Heydt et al., 2009); and *Candida orthopsilosis*, an opportunistic fungus known to cause infections in immunosuppressed humans (Riccombeni et al., 2012).

### 4.5 Future directions

The detection of RNA viruses and *Nosema* spp. spores are important for honey bee disease monitoring, but they require a different protocol from the ones presented here. The majority of viruses known to affect honey bees thus far are single-stranded positive-sense RNA viruses (Grozinger & Flenniken, 2019): however, very little is known regarding double-stranded RNA viruses, single-stranded negative-stranded RNA viruses and single-stranded RNA viruses using reverse transcriptase in their replication strategies (Payne, 2017). The implementation of eRNA will allow researchers not only to diagnose multiple known honey bee viruses simultaneously, but also speed virus discovery, advancing the honey bee virology field.

In some cases, *Apis* sp. may be one of the targets of eDNA sampling and monitoring, e.g., to detect the presence of *Apis cerana*, other *Apis* spp., or an unwanted *Apis mellifera* subspecies. However, neither region of COI nor 16S targeted by the primers we used are optimal for this application. Development of primers specific to this application, i.e., that target informative gene regions, are recommended. In contrast, there are times when honey bee DNA is not needed/targeted. In this case, one may want to “block” *Apis* DNA using primers that do not amplify it or by removing host-derived genes from the dataset. However, this step would optimally be done before PCR-amplification/enrichment, as most of the amplified DNA is likely to be *Apis* for most sample types.

We have provided evidence that collecting DNA from hive components and environmental samples in an apiary/surrounding the apiary provides robust information on organisms that honey bees contact. This, in turn, translates into an objective overview of pest presence/absence and could be modified to determine *A. mellifera* subspecies identity (e.g., honey bees of African descent), emerging pests/pathogens, etc. – all without sampling live bees. We tested and assessed efficacy and accuracy of sampling protocols enabling the timely detection of pests and pathogens correlated with honey bee health, as well as other organisms honey bees contact. These sampling protocols can be implemented without the need for advanced laboratory skills. Ultimately, the methods described in this work constitute a comprehensive molecular predictor tool of colony health. Furthermore, they can provide a timely detection of ubiquitous and/or cryptic organisms, namely pests and pathogens, before the onset of disease or before heavy pests loads occur. This could help beekeepers avoid costly treatments and negative economic consequences to their commercial enterprises.

## Data Archiving Statement

Upon acceptance for publication, raw data will be archived on Dryad.

### Acknowledgements

We would like to thank Randy Fernandez (Communication Specialist, Entomology and Nematology Department, University of Florida, Gainesville, FL, USA) for generating image graphics, David Moraga (NextGen DNA Sequencing Scientific Director, UF|ICBR) for technical assistance and Cameron Zuck (Entomology and Nematology Department, University of Florida, Gainesville, FL, USA) and Wendy Wilber (Center for Landscape Conservation & Ecology, University of Florida, Gainesville, FL) for field support. This work was supported financially by USDA-APHIS-PPQ AP20PPQS&T00C047 and the USDA National Institute of Food and Agriculture, Multi-State Hatch Project 1019945.

## Author contributions

LB conceived the study. LB, REV, HB, JDE designed the study and obtained funding. JAPM, REV and JS contributed to the acquisition of the data. Data were analyzed by REV and interpreted by all co-authors. LB and JAPM wrote the first draft, which was reviewed, edited, and approved by all co-authors.

## Supporting information

**Table S1.** Detailed sample information. Project sample dataset listing item sampled, sample collection method and environment housing the item. Visual inspection of hive health status and DNA yields per item sampled are included. (Table S1 - Detailed sample information.xlsx)

**Table S2.** Primer information and plates design. (Table S2 – Primer plates design.xlsx)

**Table S3.** Pairwise assessment of dispersion between different sample types for each primer set. Numbers indicate adjusted P-values. Significance (P<0.05) is shown in red. Where no number is provided, the pairwise comparison could not be made for that data. The sample abbreviations are defined in Table 1. (Table S3 – Dispersion results.xlsx)

**Table S4.** Amplicon metadata with taxonomic assignments for arthropods (COI and 16S), bacteria (16S) and fungi (ITS). (Table S4 – Amplicon metadata.xlsx)

**Figure S1.**
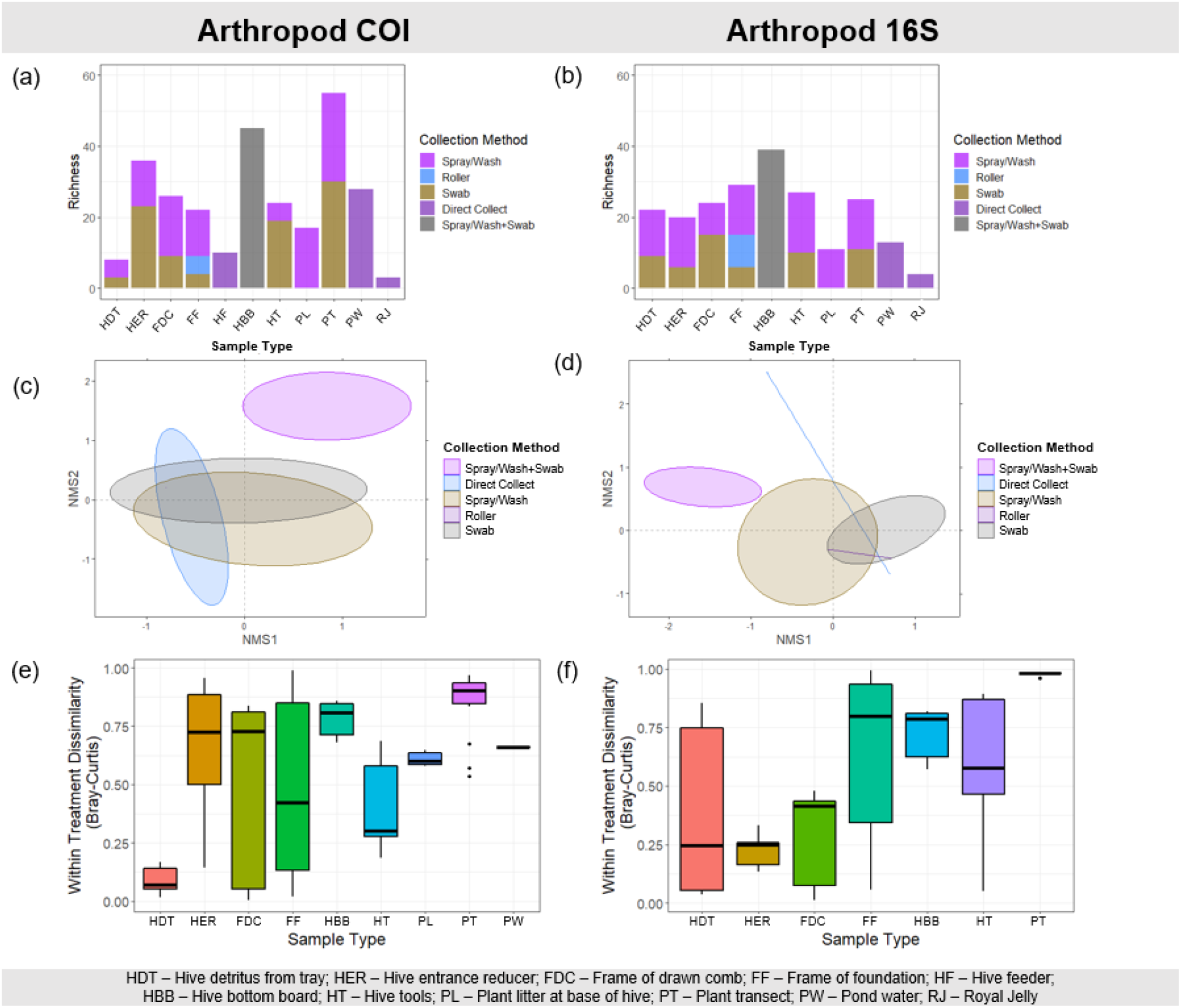
Arthropod sample type and collection method results. Richness per sample type and collection method for COI (a) and 16S (b) primer sets. Ordination plots show (dis)similarities between the different collection methods (c, d), while dissimilarity plots show (dis)similarities between replicates for each sample type (e, f) and each primer set.

**Figure S2.**
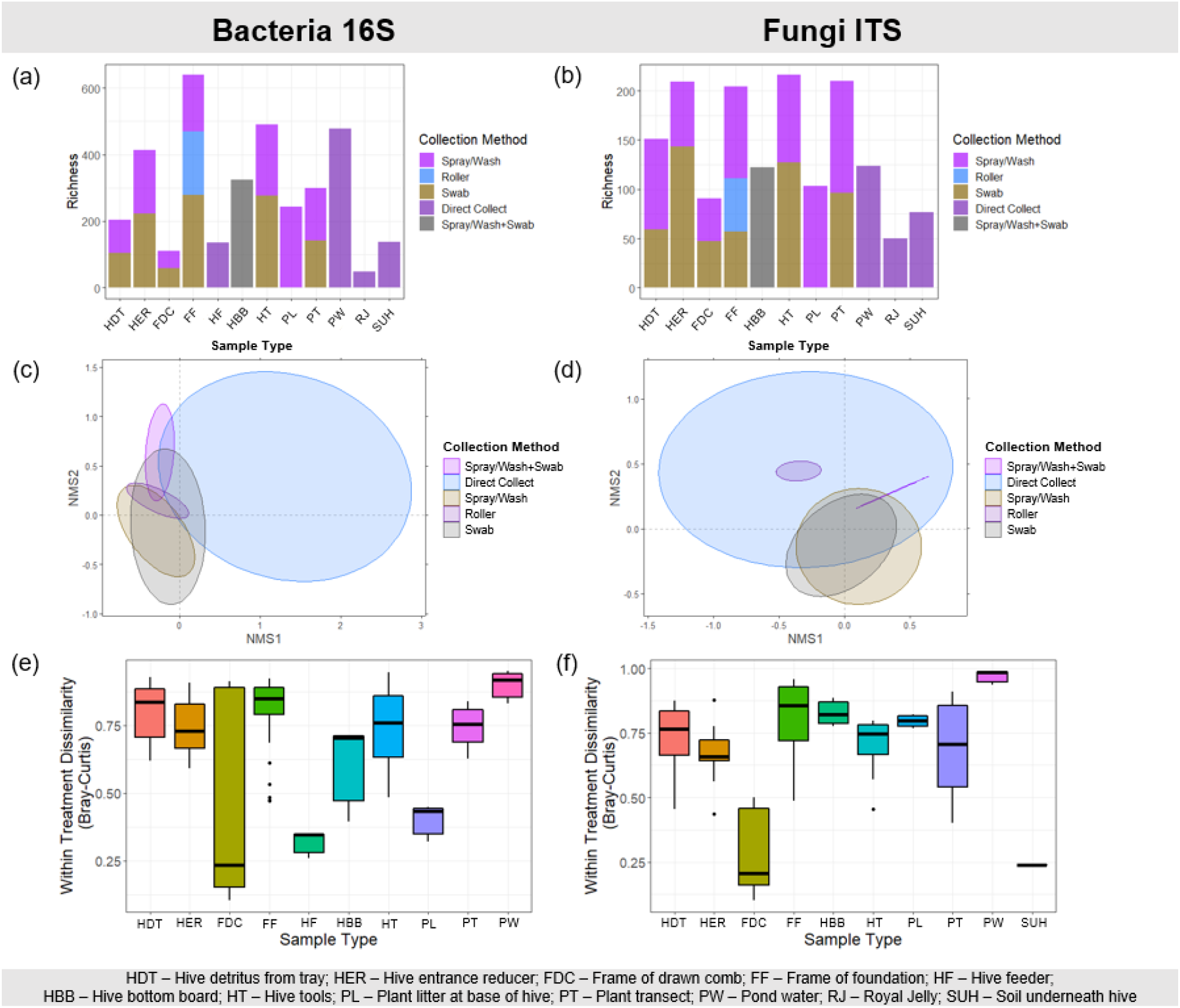
Microbial sample type and collection method results. Richness per sample type and collection method for bacterial 16S (a) and fungal ITS (b) primer sets. Ordination plots show (dis)similarities between the different collection methods (c, d), while dissimilarity plots show (dis)similarities between replicates for each sample type (e, f) and for each primer set.

**Figure S3.**
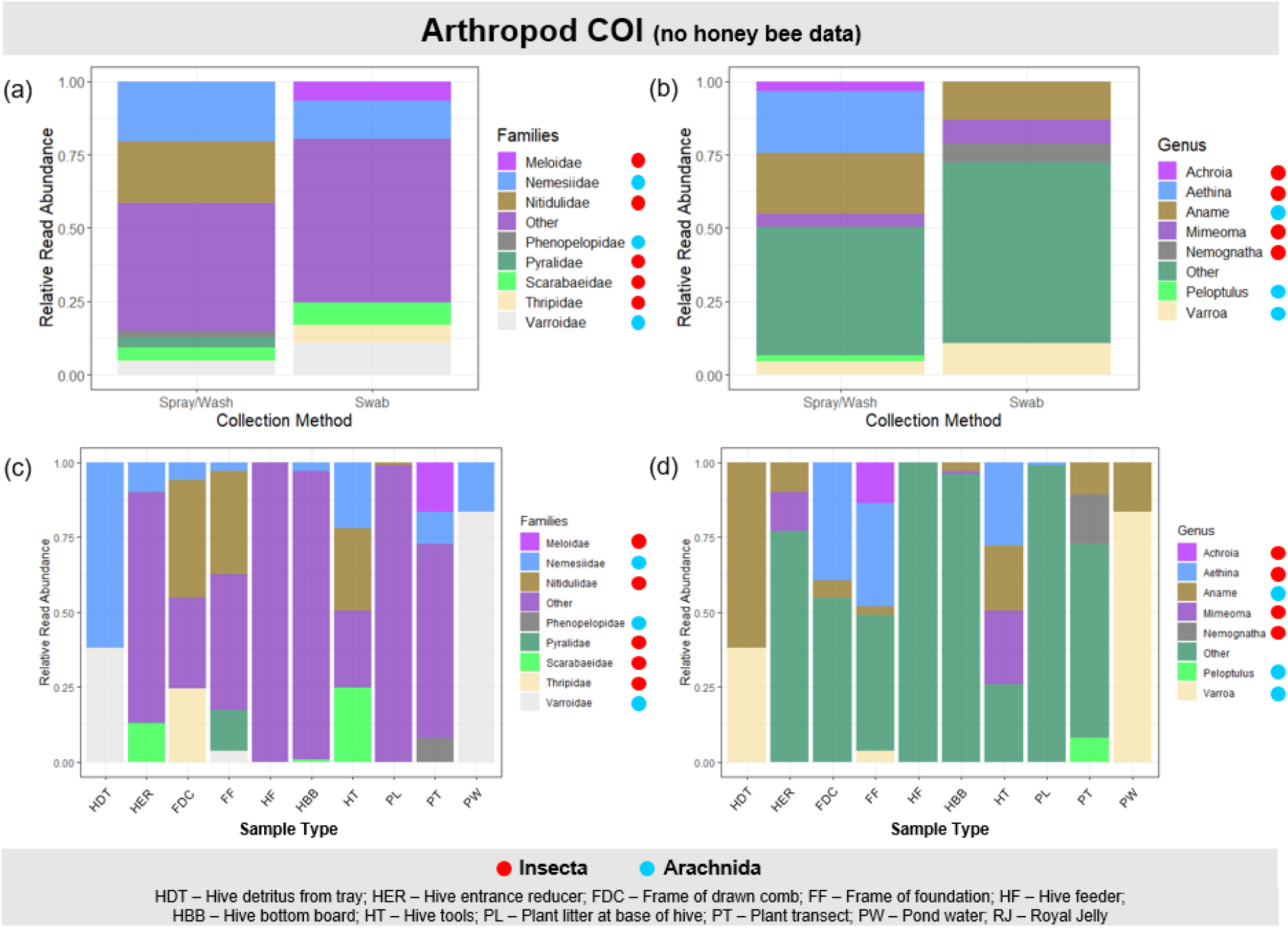
Results of the top 30 molecular Operational Taxonomic Units (mOTU) representing 83.94% of the mOTUs for the arthropod COI region, excluding host DNA amplicon reads, i.e., *Apis mellifera*. Relative Read Abundances of mOTUs are shown at the Family (a, c) and Genus (b, d) levels and are separated per collection method (a, b) and sample type (c, d). Samples not shown were discarded during the bioinformatic pipeline.

**Figure S4.**
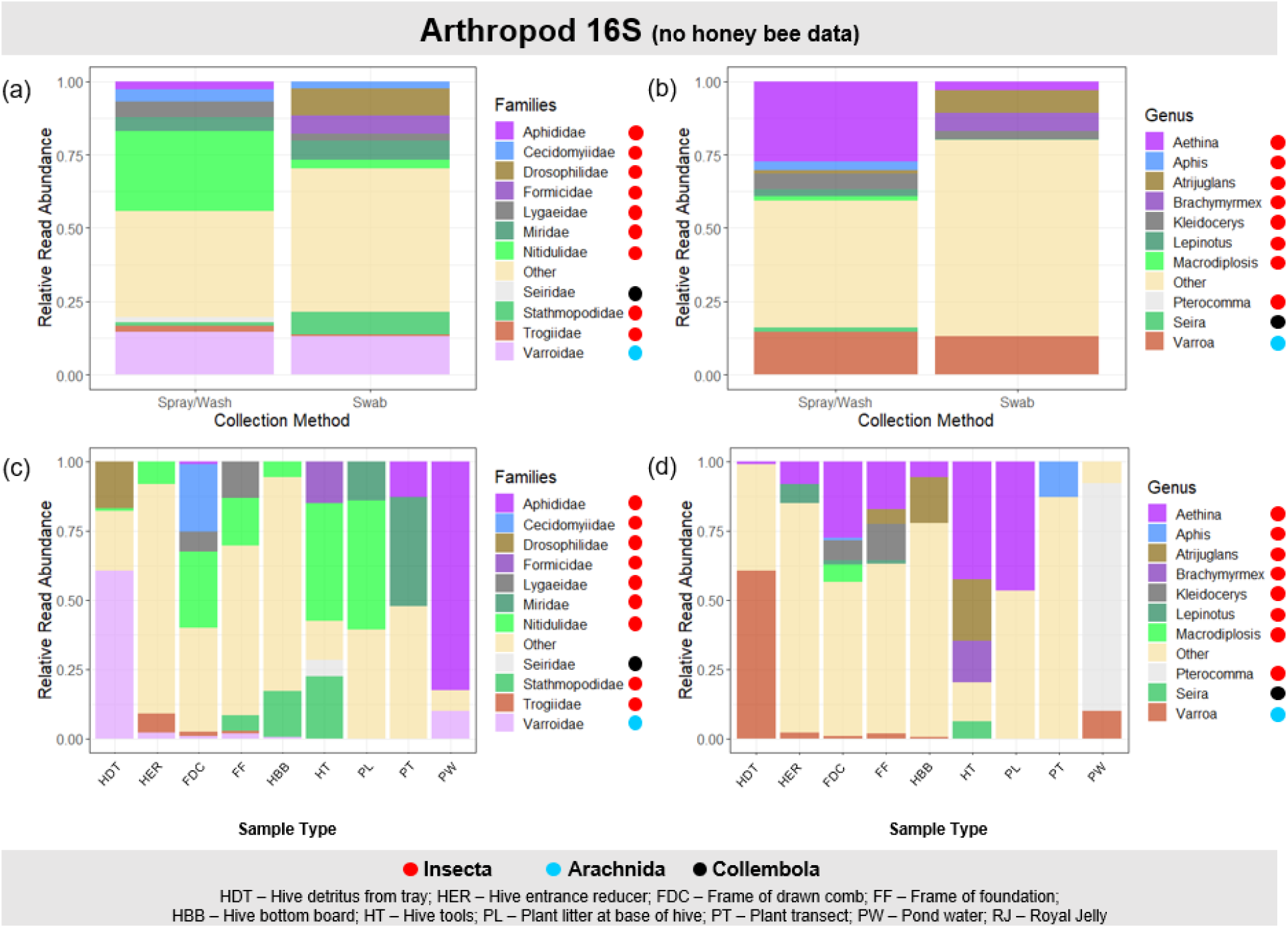
Results of the top 42 molecular Operational Taxonomic Units (mOTU) representing 96.47% of the mOTUs for the arthropod COI region, excluding host DNA amplicon reads, i.e., *Apis mellifera*. Relative Read Abundances of mOTUs are shown at the Family (a, c) and Genus (b, d) levels and are separated per collection method (a, b) and sample type (c, d). Samples not shown were discarded during the bioinformatic pipeline.

